# SCN^-^ induces prestin contraction through reduced intermonomer angles and a modified transporter like movement of each monomer

**DOI:** 10.64898/2026.10.28.762595

**Authors:** Richard Mariadasse, Molly Powers, Jun-Ping Bai, Chenou Zhang, Sneha Subramaniya, Jie Yang, Fred J. Sigworth, Venkat Kalyana Sundaram, Carmen Butan, Oliver Beckstein, Joseph Santos-Sacchi, Dhasakumar Navaratnam

## Abstract

Prestin is the membrane based molecular motor responsible for outer hair cell electromotility that expands and contracts in response to transmembrane voltage. How voltage brings these about is unknown but cryoEM structures in the presence of ligands has led to the extrapolation of structures with the transporter cycle and its expansion contraction coupling. We find prestin to adopt novel increasingly contracted states that are brought about by a reduced intermonomer angles in the obligate dimers and a transmembrane contraction of each monomer at its cytoplasmic half in the presence of SCN^-^. These changes are associated with a loss in the intermonomer beta sheet at its N terminus. There is a hyperpolarizing shift in *V_h_* along with increased affinity for SCN^-^. SCN^-^ binding at the internal binding site likely cause the structural changes since they are reversed by SCN^-^ displacement by salicylate.

## Introduction

Prestin (SLC26A5) is the molecular motor responsible for electromotility in outer hair cells that in turn brings about cochlear amplification (*1–5*). It is a member of the SLC26A family of anion transporters that has 10 members (*6*). To date, cryoEM structures have been determined in eight of them, that reveal a consistent dimer with an intracellular N and C terminus that bookend an inverted repeat 14 transmembrane region divided into a core domain that has been modeled to move in an elevator like motion on a gate domain (*7–12, 13, 14*). Of 7 structures (we excluded SLC26A11 that differs from the others (*14*) and will not be considered here), all but pendrin (Slc26a4) is in an inward facing or inward occluded state. Pendrin structures were found to be in an inward facing, inward and outward facing and outward facing structure using a mixture of substrates (Cl^-^/ HCO3^-^) that contrasts with the uniform structures that were obtained with mostly single substrates for the other cryoEM structures (*10*). Prestin is a motor protein that responds to transmembrane voltage expanding and contracting with hyperpolarizing and depolarizing voltages (*3, 15, 16*). The cryoEM structures of prestin reveal a mostly inward occluded state and show some variation with different ligands that have been extrapolated to changes in voltage (*11–13*). In this paper, we use a mixture of SCN^-^, that elicits robust currents with prestin(*17, 18*), and Cl^-^ and find several prestin cryoEM structures revealing a more contracted protein brought about by closer approximation of each monomer and an inward movement of the transmembrane region mostly at its cytoplasmic half.

## Results

### In the presence of SCN^-^, prestin takes on increasingly rotated and somewhat outward facing structure

We determined the cryoEM structure of prestin in the presence of 40mM SCN^-^ and 100mM Cl^-^. We identified four structures that overall maintained the core dimeric structures previously observed in the presence of Cl^-^ and SO ^2-^. Figure 1 shows the density map and ribbon diagrams of these structures and confirms their intracellular N terminus, the 14 transmembrane regions containing the gate and core domains and the intracellular C terminus with its STAS domain. The structures, 9YAY, RM1, 2 and 3 had a resolution of 3.8A, 3.56A, 3.88A, 3.65A, respectively (using the gold standard FSC). One structure (9YAY), most closely resembled three previous structures of prestin in Cl^-^ (7SUN, , 7 LGU and 7s8x with RMSDs in the Tm domains of 0.571, 0.451, and 0.477A, respectively (*11, 12, 19*)) in an inward occluded state similar to Slc26a9 (overall RMSD 3.2 (*20*)). We have already referenced the 9YAY structure in a related paper, where we identified the presence of an external anion binding site (*21*). One structure, RM2, more closely resembled the recently described outward facing structure of pendrin (Figure 1). Two other structures (RM1 and 3) took on conformations that were intermediate between 9YAY and RM2 (supplementary Figure 1). We have included 9YAY for consistency and in order to make valid comparisons since it too was obtained in the presence of SCN^-^. Figure 2 shows the 9YAY and RM2 structures where we have aligned the gate domains of monomer A with that of monomer A of the inward facing pendrin structure (7wlk1) and shows the relative rotational and upward movement of prestins core domain (*10*). Included are the outward facing structure of pendrin that was similarly aligned. The figures show the relationships of the Tm domains 3 and 10, that forms the critical internal Cl^-^ binding site in the family, undergoing a progressive rotational and modest upward deviation in RM2. Also evident are the Tm regions of the core domain and its similar rotational and upward movement compared to the inward facing pendrin (7wlk1) (*10*). Tm1 shows the greatest movement. Supplementary table 1 shows the RMSD values of the core and gate domains and individual Tm domains when the structures were aligned with the gate domain of 7wlk1. Figure 2 also shows a relative alignment of the gate domain of RM2 with 7wlk1 but for the C terminal end of Tm6 that shows a significant deviation. These data extend Bavi et als findings of a hinge region in Tm6 (*11*).

**Figure 1.**
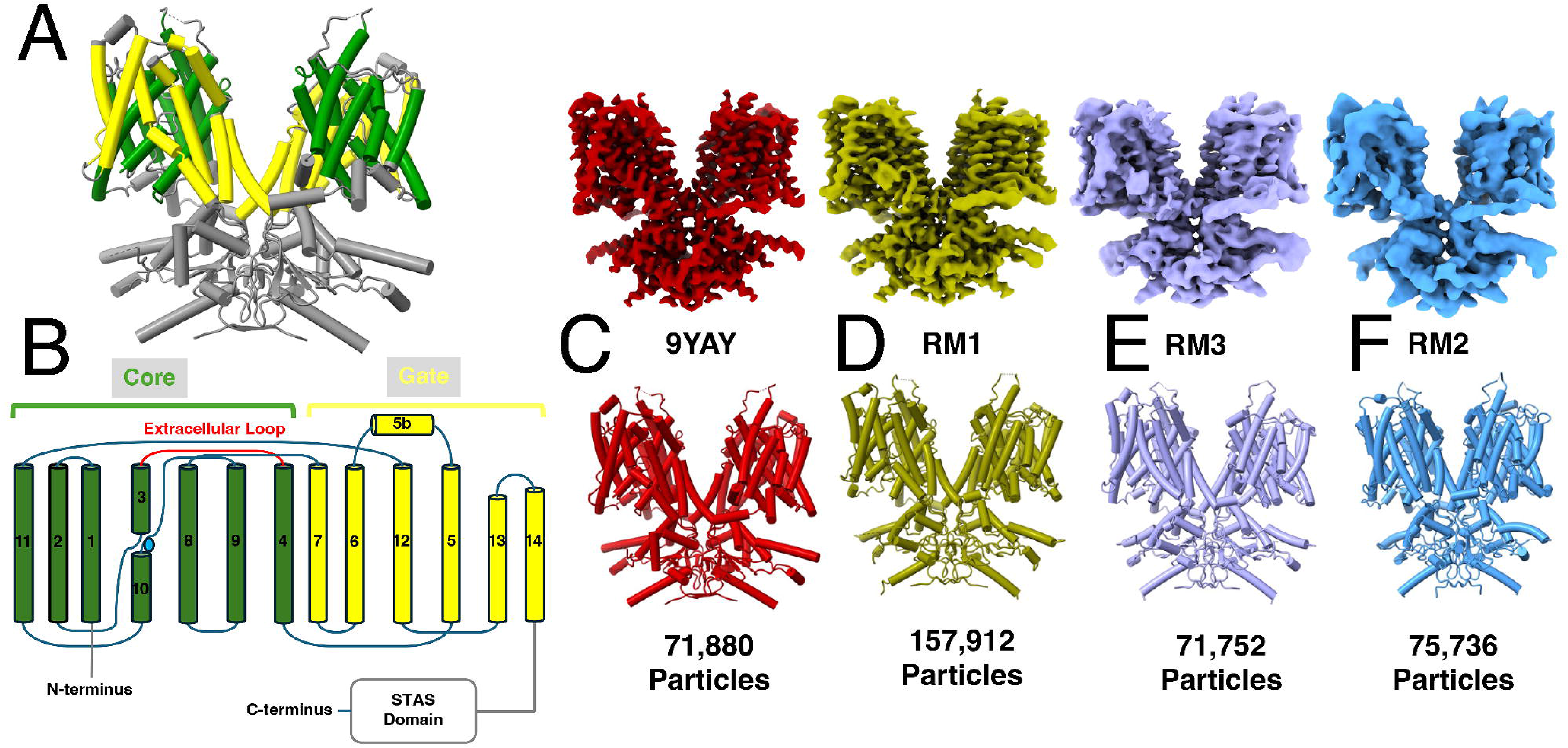
Four prestin structures are identified in the presence of 40mM SCN^-^. **(A,B)** Shown are the atomic model of prestin dimers and a cartoon of the monomer showing the core (yellow) and gate (green) transmembrane domains along with the intracellular C and N termini. **(C-F)** Shown are the electron density maps (upper) and respective atomic models (lower) of the four structures 9YAY (C), RM1 (D), RM3 (E) and RM2 (F) including the number of particles that were used to derive the structures.

**Figure 2.**
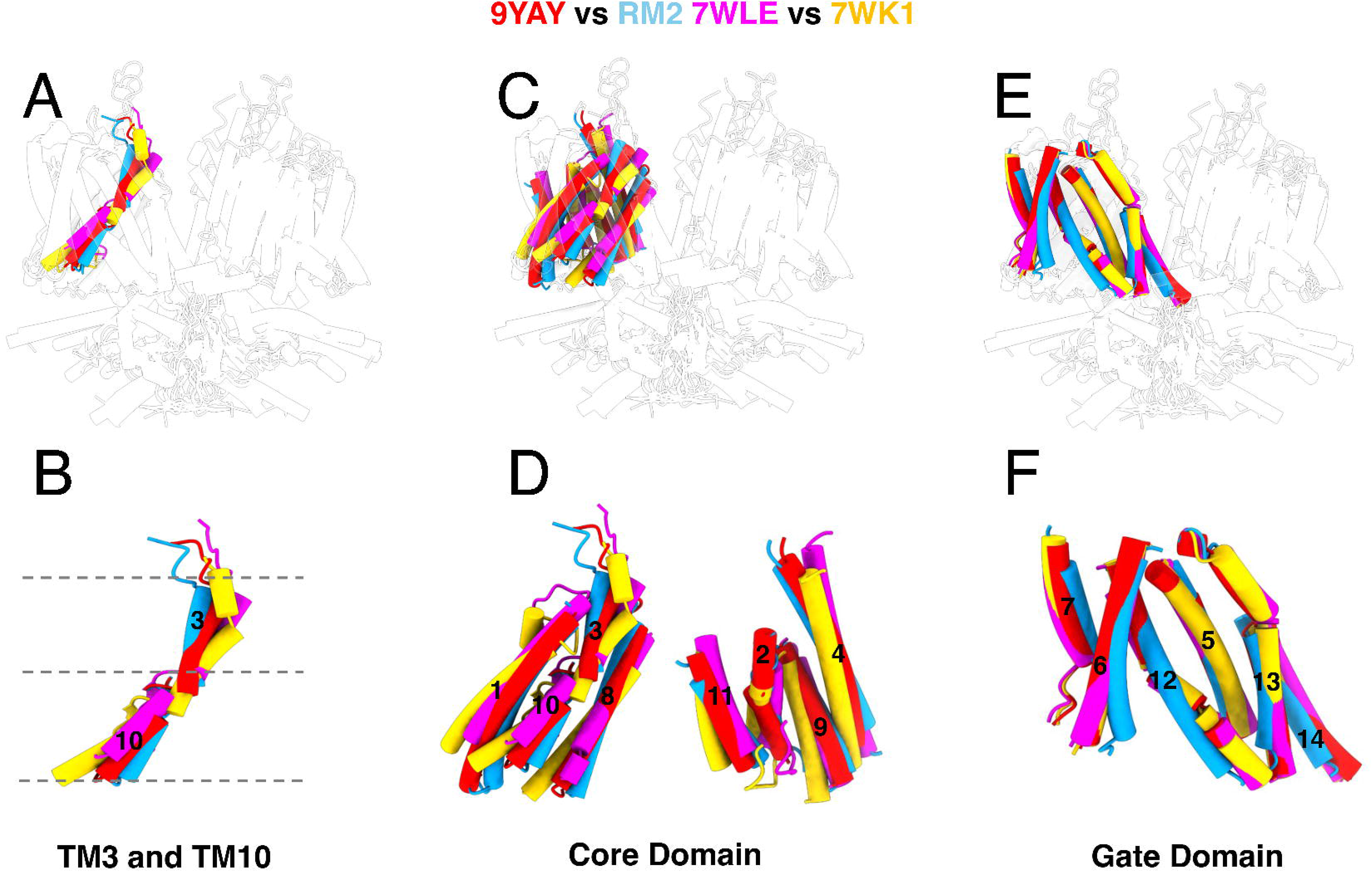
There is a progressive rotational and upward movement of the core domain on the gate domain in RM2 when compared to the inward (WK1) and outward (WLE) facing pendrin structures. The gate domains of one monomer of the four structures and the outward facing pendrin structure (WLE) were aligned with the inward facing pendrin structure (WK1). **A-B.** Show the location of Tm domains 3 and 10 within the structure of the dimer **(A)** and the relative movements of Tm3 and10 of RM2 and 9YAY when compared to the inward and outward facing pendrin structures **(B)**. The progressive rotational and upward movements from 9YAY to RM2 of Tm3 and 10 from 7Wlk1 to 7Wle is evident. **C-D.** Show the rotational and upward movements of the inner four (Tm 1, 3,10, and 8) and outer four (Tm 11, 2, 9 and 4) Tm segments of the gate domain from 9YAY to RM2 compared to 7Wle and 7Wk1. **(E-F)** Show the gate domains of the four structures and 7Wle aligned to 7Wk1. RM2 shows a deviation of the most cytoplasmic half of Tm6.

### The rotational and upward displacement of the core domain correlates with a decreasing angle between the two protomers and a contraction in the transmembrane region at the inner leaflet

We note reduction in the distances between the two protomers in RM2 evidenced by a decrease in the angles (ø) between the axes of the transmembrane regions of the two protomers (here we used the principal component axis of the transmembrane region from aa 75-505 of each protomer). RM1 and 3 show a similar but lesser reduction in the distances and angles. This is shown in Figure 3. ø decreased from 69^0^ in 9YAY to 64^0^, 61 and 33^0^ in RM1, 3 and 2 respectively. For comparison, outward facing pendrin too shows such a reduction that is, however, smaller (ø is 45^0^ in IF and 34^0^ in OF respectively). Interestingly, the changes in ø does not correspond to a change in the angle of axis between the Tm and intracellular C terminus that remains at ∼ 140^0^ (9YAY:141, RM1 141, and RM3 141) except in the most contracted form (RM2) where it changes to 133^0^. These reductions in the angle between the protein also cause a reduction in the area between the protomers. Coincident with the reduced angles between the protomers, we observe the absence of the beta sheet between the N termini of the two protomers (Figure 2). All three prestin structures that show a reduction in the angles between the protomers have this feature while the structure in the presence of SCN^-^ (9YAY) that most closely resembles the inward occluded structure (7SUN) and maintains a normal interprotomer angle retains its beta sheet. While we do not know if this absence of the beta sheet is causally related, we note that deletion of the N terminus is linked to a loss FRET signal between monomers (*22*).

**Figure 3.**
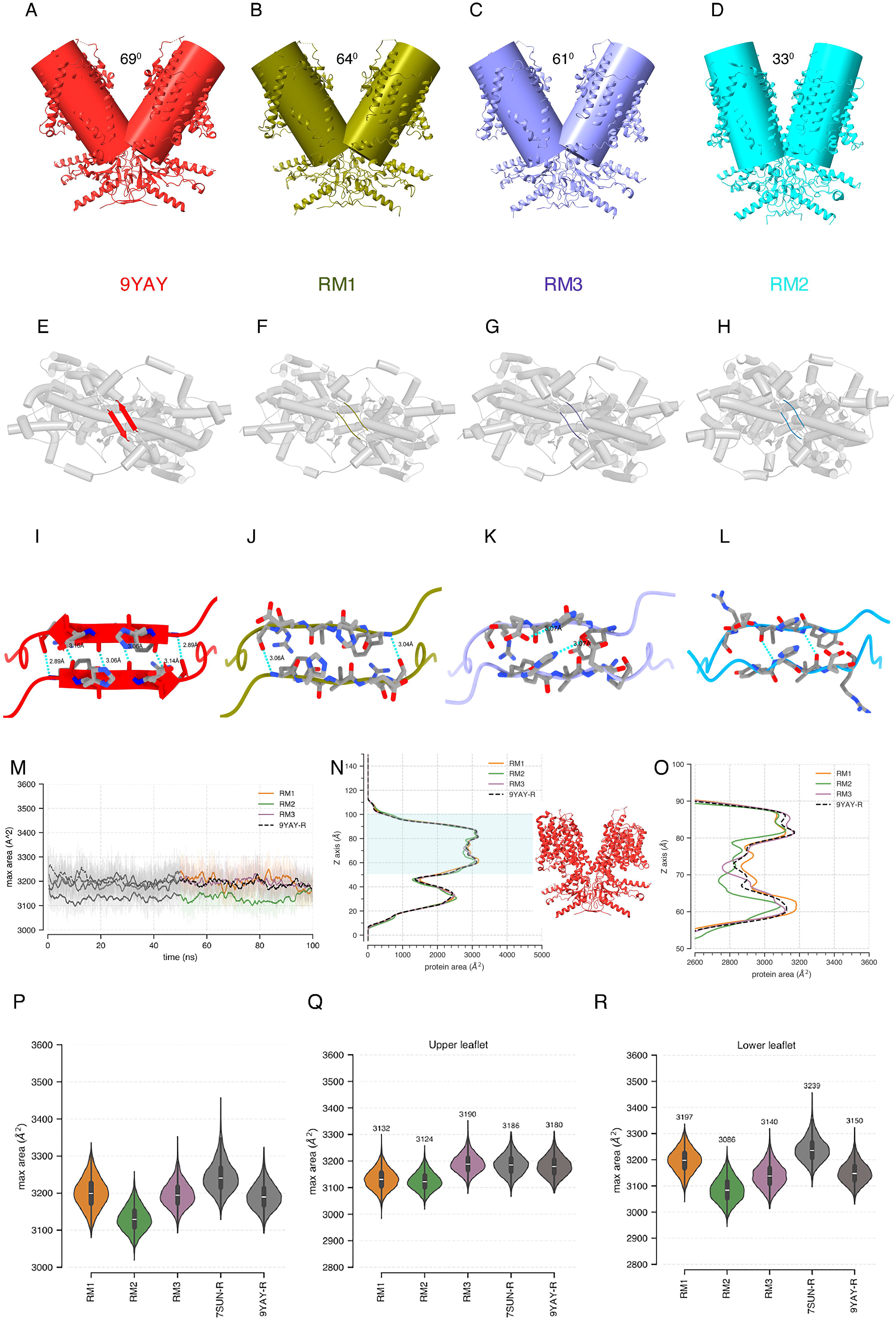
There is a progressive reduction in the angle (ø) between the protomers that is parallelled by a reduction in the area of each dimer. **A-D** Shows a progressive reduction in the angles between the monomer pairs of each dimer in the presence of SCN^-^. This is associated with a loss in the short beta sheet formed at the N terminus between monomers **(E-L). I-L.** Shows the side chain interactions along with their H bonds. Most of the six H bonds that form the beta sheet in 9YAY are disrupted in RM1-3. **M-R.** Shows the reduction in area of each of the dimers in the presence of SCN^-^ compared to Cl^-^. **M-O.** Shows a restrained simulation of the structures over 100ns showing the maximum area of the protein over the entire z coordinate at 10pS intervals. (**M).** The time-averaged protein area of the protein was then plotted as a function of the z coordinate. The system was allowed to equilibrate for ∼the first 50ns and data from the last 50ns used for statistical comparisons between the different structures. **(N-O).** The membrane area is shaded in **N** and the Tm segments of the model abutted. **O** is an expanded view of **N**, showing the reduced area most notably in RM2. **P-R** are violin plots of the areas of each simulation demarcating the maximum area of the entire transmembrane area **(P)**, and maximum area of closest to the outer **(Q)** and inner leaflet **(R)** respectively of each of the structures showing a progressive reduction in area of all these structures compared to prestin (7SUN) in the presence of Cl^-^. As is evident the contraction of the protein is asymmetric and mostly at its inner leaflet.

We also sought to determine if the differing structures showed differences in transmembrane area. To do so we used a restrained Molecular Dynamics simulation to ascertain the relative area change in the protein. Figure 3 shows the area changes of RM1-3 compared to 9YAY and 7SUN, the conformation in the presence of Cl^-^. As is evident, all the structures in the presence of SCN^-^show a reduction in area compared to that in Cl^-^. Moreover, there is a progressive reduction in the areas of these four structures with the area of the most contracted dimer (RM2) being ∼65Å^2^ less than that of 9YAY and 113 Å^2^ less than 7SUN. Figure 3 also shows the distribution in area across the Tm domain and confirms that the contraction of the protein across the membrane is not uniform but rather concentrated closer to its inner leaflet, similar to our modeling data (*21*). This is most evident in Tm1 and Tm2 that shows the most movement (see supplementary Table1 for RMSDs of the different Tm domains compared to 7SUN). These findings are consistent and confirm our simulations of the protein in the presence of Cl^-^ (*21*). As previously noted by Bavi et al., the segment closest to the inner leaflet of Tm6 also shows significant movement that contributes to area reduction (in their case expansion with SO ^2-^ and salicylate but also see below (*11*)) (*10*).

### The rotational and upward displacement of the core domain correlates with a decreasing angle between the two protomers and a contraction in the transmembrane region at the inner leaflet

We note reduction in the distances between the two protomers in RM2 evidenced by a decrease in the angles between the axes of the two protomers. RM1 and 3 show a similar but lesser reduction in the angle. This is shown in Figure 3. The angles decreased from 69^0^ in 9YAY to 64^0^, 61^0^ and 33^0^ in RM1,3 and 2 respectively. Interestingly, the changes in the angle between prestin protomers does not correspond to a change in the angle of axis between the Tm and intracellular C terminus that remains at ∼ 140^0^, (9YAY:141.35, RM1 141.5, and RM2 141.1) except in the most contracted form where it changes to 133^0^. These reductions in the angle between the protein also cause a reduction in the area between the protomers. Coincident with the reduced angles between the structure, we observe the disruption of the beta sheet between the N termini of the two protomers (Figure 3, and supplementary Figure 2). All three prestin structures that show a reduction in the angles between the protomers have this feature while the structure in the presence of SCN^-^ (9YAY) that most closely resembles the intermediate inside open structure (7SUN) retains its beta sheet. While we do not know if this absence of the beta sheet is causally related, we note that sequential deletion of these residues at the N terminus is linked to a loss FRET signal between monomers (*22*). Importantly, the pendrin OF structure that shows a reduction in its angle also has a loss of this beta sheet observed in the IF structure (*10*).

We also sought to determine if the differing structures showed differences in transmembrane area. To do so we used a restrained molecular dynamics simulation to ascertain the relative area change in the protein. Figure 3 shows the area changes of 9YAY and RM1-3 compared to 7SUN. As is evident, there is a progressive reduction in the areas of these three structures with the area of the most contracted dimer (RM2) being ∼113 A^2^ less than that of 9YAY. Figure 3 also shows the distribution in area across the membrane and confirms that the contraction of the protein across the membrane is not uniform but rather concentrated closer to its inner leaflet, similar to our previous modeling data (*21*). This is most evident in Tm1 that shows the most movement (see supplementary Table1 for RMSDs of the different Tm domains compared to 7SUN.

### SCN^-^ binds to prestin with a higher affinity at the internal binding site and causes a hyperpolarizing shift in *V_h_*

Why does SCN^-^ cause a contracted and outward facing structure? Figure 4 shows that prestins voltage of peak non-linear capacitance (NLC, that is a surrogate measure for electromotility) shifted in a hyperpolarizing direction in a range of SCN^-^ concentrations compared to Cl^-^, while other measures of charge movement including *Q_sp_*, the charge movement per unit of membrane, and z, a measure of the charge movement of a single motor subunit, were indistinguishable from that in the presence of Cl^-^ (supplementary Figure 3). This shift in *V_h_* could be the reason we see a more contracted /compact form in the presence of SCN^-^, as detergent micelles don’t have a volage gradient across them and are effectively at 0mV. Since there is no change in z, these findings argue that the contracted structures we observe are normative and likely represent the full measure of contraction not captured in the previous structures in the presence of Cl^-^.

**Figure 4.**
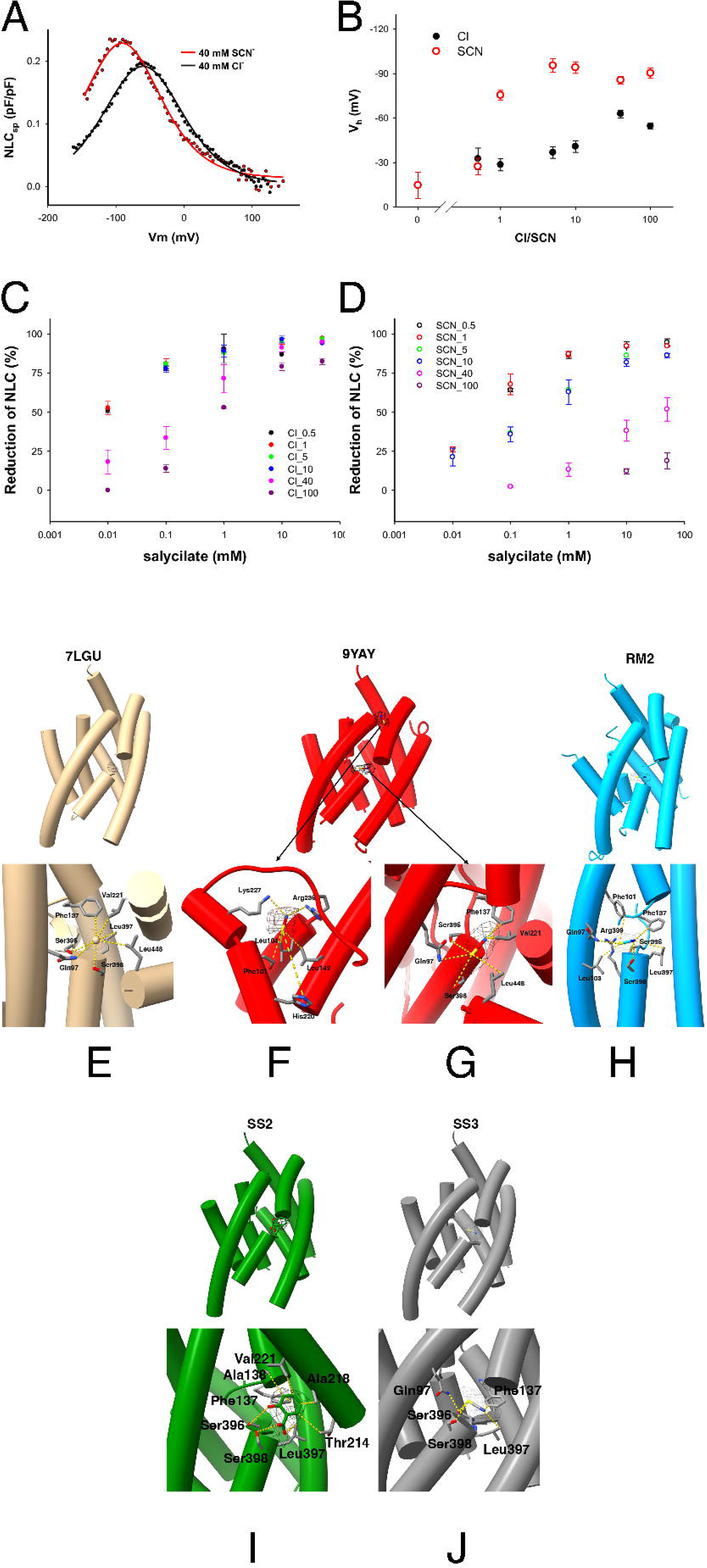
SCN□ causes a hyperpolarizing shift in prestin’s voltage dependent NLC response and reduces salicylate inhibition, likely from higher affinity to the internal binding site. NLC was recorded from a prestin stable cell line with intracellular Cl□ or SCN□, in the absence or presence of salicylate. **(A)** Representative NLC traces with 40 mM intracellular Cl□ or SCN□, illustrating an SCN□-dependent shift in *V_h_*. **(B)** NLC recorded with intracellular Cl□ or SCN□ at the indicated concentrations was fit with a two-state Boltzmann function to obtain the midpoint voltage *V_h_*; SCN□ produces a concentration-dependent depolarizing shift in *V_h_*relative to Cl□. **(C, D)** Effect of salicylate on NLC: salicylate nearly abolishes NLC in all concentrations of Cl□, consistent with displacement of bound anion from the internal binding site, whereas substantial NLC persists in SCN□ even at the lower SCN□ concentration tested, indicating incomplete displacement and suggesting higher affinity/competition by SCN□. **E-H.** Shown are atomic models of prestin with Cl- (Ge et al., 2021) at the internal binding site of 7LGU **(E)**, and SCN- at the internal binding site of 9YAY **(F)** and RM2 **(G)**. Cl^-^ forms polar bonds with S398OG, S398 backbone N, S396, L397 backbone, and Q97 while the S atom of SCN forms similar polar bonds with these same residues **(G)**. In addition, the N atom of SCN forms nonpolar bonds with F137 and V221.

What could cause the hyperpolarizing shift in *V_h_*? Since Cl^-^ binds to the internal binding site, we reasoned that SCN^-^ does the same and tested if salicylate that displaces Cl^-^ from the internal binding site could similarly displace SCN^-^ from the internal binding site. Figure 4 shows the effects of different concentrations of SCN^-^ and salicylate on NLC measures and contrasts it with the effects on Cl^-^. Salicylate displaces Cl^-^ with a near complete loss in NLC even at a 100mM Cl^-^. In contrast, salicylate is unable to completely reduce NLC in the presence of SCN^-^. Thus, at similar concentrations of anions, salicylate has reduced ability to inhibit NLC in the presence of SCN^-^ suggesting that SCN^-^ has a higher binding affinity.

What could account for the higher affinity of SCN^-^ to the internal binding site? Figure 4 shows the coordinating residues that bind SCN^-^ in 9YAY and compares it to Cl^-^ in 7LGU, the structure that is most close to 9YAY. In 7LGU, the radial distribution function has a sharp first peak at 4.38Å and a first minimum at 5.46 Å with bonds that are largely polar (S398OG 3.54 Å, S398 backbone N 3.61 Å, S396 OG 3.66 Å, L397 backbone N 3.7 Å, and Q97 sidechain N 4.11 Å). A second peak at 6.48 Å with a minimum of 7.68 Å was not considered further. In the case of SCN^-^, the S atom has an inner shell with a peak at 4.22 Å and polar bonds (Q97 OE1 3.49 Å, Q97 side chain N 3.88 Å, S398 backbone N 4.02 Å, S396 OG 4.08 Å and S398 OG 4.51 Å). A second shell peaks at 5.3 Å. The N atom of SCN^-^ interacts with a hydrophobic pocket with the first peak RDF of 3.2 Å (F137 (CE1 3.05 Å, CD1 3.24 Å), V221 (CG2 3.44 Å)) and a second peak at 5.3 Å. Thus, the two ends of SCN sit in different environments. The S end occupies the polar part of the site: Q97, the S396 and S398 hydroxyls, and the S398 backbone amide. These are largely the same groups that hold chloride in 7LGU. The N end projects into the hydrophobic sub-pocket formed by F137, V221, L397 and F101. We reason that the latter interactions give SCN^-^ its enhanced binding.

### Salicylate displaces SCN^-^ from the internal binding site and restores prestins structure closer to 7SUN along with restoration of the angle between the two protomers

In order to confirm that the structural changes in prestin were due to the binding of SCN^-^ at the internal binding site, we determined structures of the protein in the presence of SCN^-^ and salicylate that competes with anions at the internal binding site (albeit incompletely with SCN^-^). Since SCN- has a higher affinity for the internal binding site compared to Cl^-^, and requires higher concentrations of salicylate, we used 10mM SCN^-^ and 10mM Salicylate to allow salicylate to compete with SCN^-^ at the internal binding site (see Figure 4 for effects on NLC in differing concentrations of SCN^-^ and salicylate). Figures 5 and 6 shows the structure of prestin (SS2) with salicylate bound to the internal binding site that takes on an expanded state and restores the angle between the monomers. The angle between the axes of monomer A and B was 69° (SS2) and similar to that of 9YAY (69°). SS2 is similar to the structure described previously (7LH2 (*13*)) in the presence of salicylate and shows a small increase in its area (+13 A^2^) along with an outward rotation and downward movement of the core domain on the gate domain compared to 9YAY. We also found two other structures in the presence of salicylate and SCN^-^(supplementary Figure 4) by 3D classification. In one structure (SS3), we find SCN^-^ still bound to the internal binding site. This structure is closest to RM1 in the presence of SCN^-^ only (RMSD of the core domain is 1.46). Importantly, the angle between the protomers is reduced at 62°. A third structure was of too low a resolution (4.8A) to identify the anion or salicylate at the internal binding site but was similar to the structure with bound salicylate (RMSD of the core domains is 1.433). Here the angle between the protomers was 70°. Notably all three structures show a beta sheet at the N terminus, although that of SS3 with bound SCN^-^ and the smallest angle between the protomers shows a significant 90^0^ turn at the very c-terminus of the beta sheet and does not show modeled interprotomer hydrogen bonds. Thus, SS3 differs from all the other structures with beta sheets at this location. Additionally, SS3 has the largest area of all the structures we determined being 82 Å^2^ larger than 9YAY. Since it has SCN^-^ at its internal binding site these data raise concerns that the expansion of the protein with salicylate may not be equivalent to the expanded state with hyperpolarizing voltage as prior electrophysiological data have shown.

**Figure 5.**
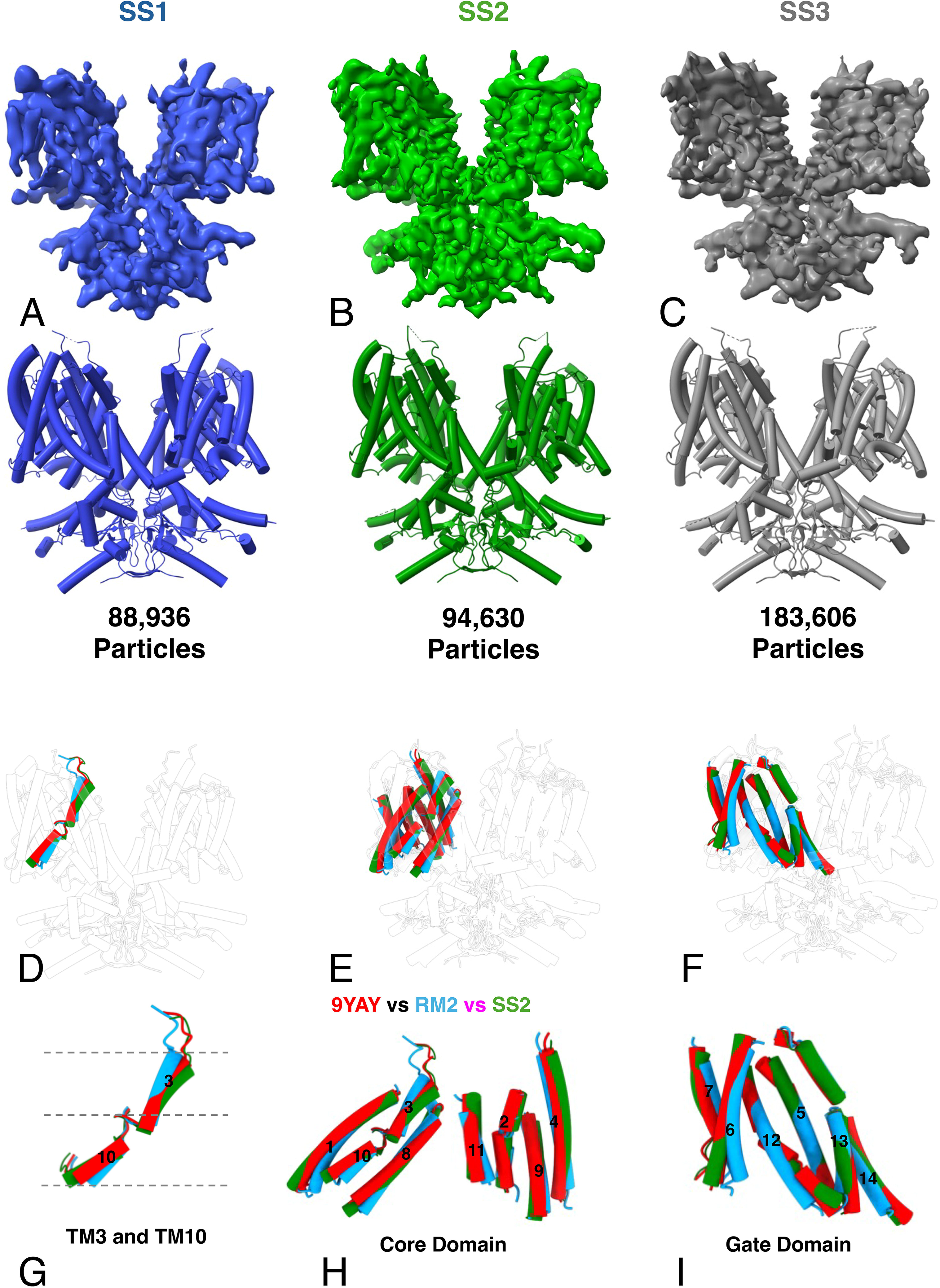
Salicylate reverses the structural changes induced by SCN^-^. **(A-C)** Shows the electron density maps (upper) and atomic models (lower) of three prestin structures in the presence of 10mM salicylate and 10mM SCN^-^ (SS1 **(A)**, SS2 **(B)**, and SS3 **(C)**). The particle numbers that were used to make these models are indicated. **(D-I)** Show the location of the Tm domains 3 and 10 forming the internal binding site **(D,E)**, the core domain **(F,G)** and the gate domain **(H,I)** after superimposing the gate domain of SS2 on the gate domain of 9YAY. SS2 has a bound salicylate ion at the internal binding site. Tm3 and10, and the gate domain superimpose on that of 9YAY, reversing the SCN^-^ induced changes in RM2. In the core domain, Tm6 also reverses its bending and aligns with 9YAY.

**Figure 6.**
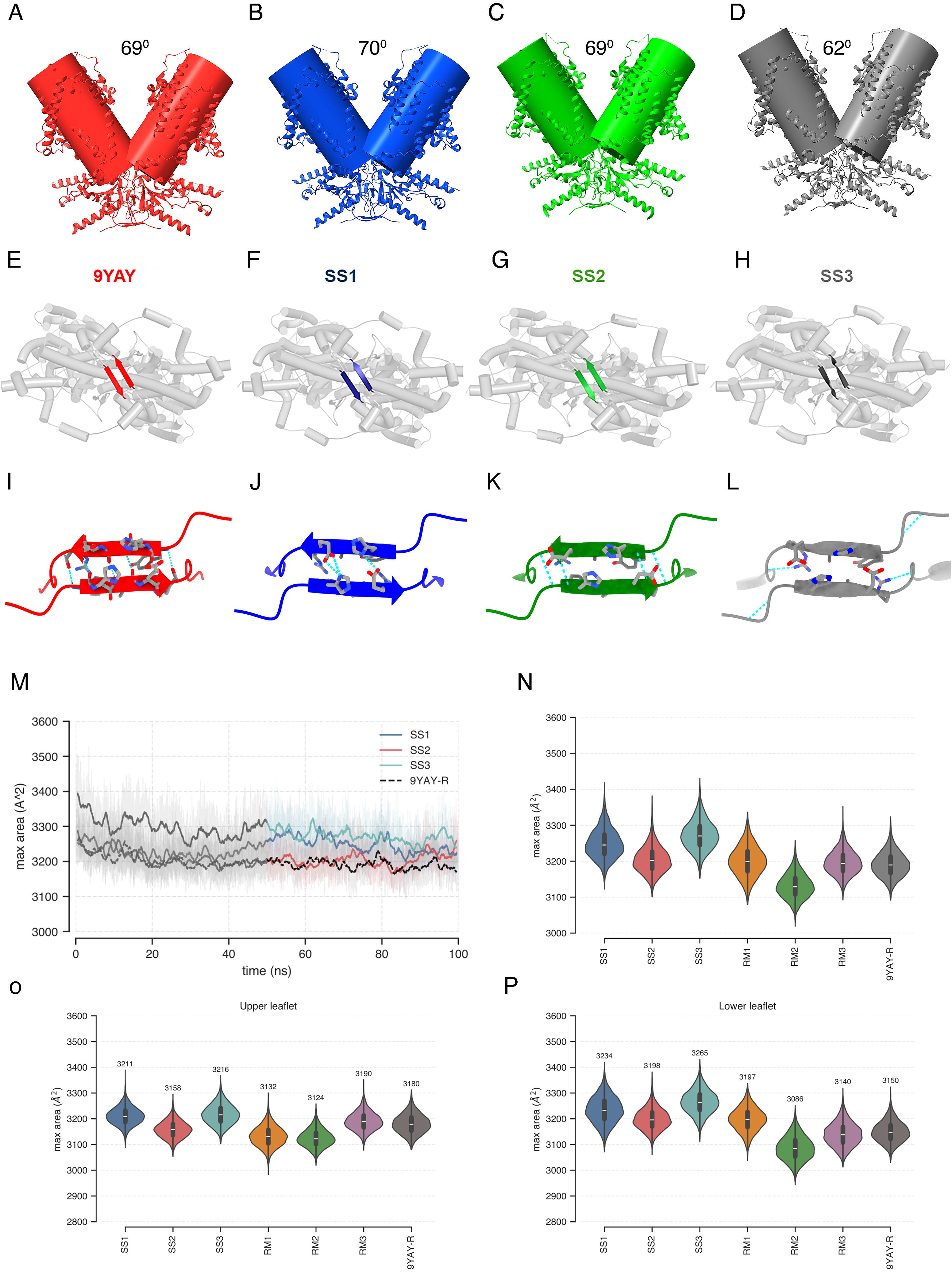
Salicylate reverses the effects of SCN^-^ on the intermonomer angle (ø) and the contracted protein. **A-D**.Shown are the angles between monomers of three structures with salicylate (SS1-3) compared to 9YAY. Except for SS3 the angles between the monomers are near identical to 9YAY. **E-H**. Shown are the presence of the beta sheet between the N terminus of each monomer. SS1 and2 show a beta sheet that is similar to 9YAY. The beta sheet of SS3, that has SCN^-^ at the internal binding site, is distorted at their C terminal ends and differs from SS1-2 and 9YAY. Note that the angle between monomers is also reduced in SS3. **I-L.** Shows an expanded view of the beta sheets and the hydrogen bonds between the two beta sheets. There are no hydrogen bonds in SS3. **M-P.** Shows restrained simulations over 100ns of each of SS1-3 **(M)** and compares the maximum area in the transmembrane region **(N)**, and the region restricted to the upper **(O)** and lower **(P)** leaflet of the membrane. As evident the structures in the presence of salicylate (SS2) have a higher area than the contracted state with SCN^-^ (RM2). The expansion is most notable in the region closer to the lower leaflet.

## Discussion

In this paper we find that prestin, in the presence of SCN^-^, adopts a conformation more close to the outward facing structure of pendrin with a notable reduction in the angle (ø) between the monomers.

The reduction in angle (ø) between the two monomers is a new finding but has parallels in pendrin where, however, the reduction on in the angle (ø) between the inward and outward facing structures is smaller. Prestin in outer hair cells is found in high density and this closer approximation of the two monomers has important implications. First, the narrowing and widening of the intermonomer angle implicates intersubunit interactions of the protein. Second, such a reduction in space would necessitate a redistribution in lipid between the monomers to an externality such as a change in the height of the membrane surface in microsecond timescales. Third, cholesterol, a lipid with significant effects on prestins NLC is found in the cleft between the two monomers and this reduction in the angle (ø) between monomers can have effects on this interaction.

The reduction in the angle (ø) between the monomers with SCN^-^ in RM2 is accompanied by a disrupted beta sheet between the N terminal ends of the monomers. The pendrin transition from inside open to outside open is accompanied by a similar reduction in angle (ø) between the monomers and, importantly, a disruption in the corresponding beta sheet at the N terminus. In the presence of salicylate at the internal binding site of prestin, the angle (ø) between the monomers and the beta sheet at the N terminus are restored, suggesting that SCN^-^ binding at the internal binding site is responsible for both of these changes. SS3 that has SCN^-^ at the internal binding site shows a smaller interprotomer angle (62^0^) and has a beta sheet at the N terminus is discordant with our conclusions. However, here, the beta sheet is twisted and lacks interprotomer hydrogen bonds in our models, indicating an unstable beta sheet. Previously, we found that mutation of these N terminal residues resulted in a loss in NLC that was accompanied by a loss in FRET signal. Our present findings lend support to the possibility that interactions at the N terminus of the two monomers stabilize the protein in an inward facing and expanded conformation. Importantly, the angle (ø) between the monomers was not significantly different between the 9YAY and SS2 suggesting that these angles (ø) had reached an asymptote. Thus, it maybe that contraction of the protein is a two-step process with intermonomer angles and contraction of each individual monomer enhancing this contraction further (*21*).

All of the structures in the presence of SCN^-^ alone that also show SCN^-^ to be present at the internal binding site have an area that is consistently less than that of prestin in the presence of Cl^-^. However, SS3, obtained in the presence of salicylate and SCN^-^, that has a clearly identifiable SCN^-^ ion at its internal binding site also has the most expanded structure. Previously, it was argued that the expanded states seen with salicylate was representative of the expansion as salicylate was thought to displace chloride and shift *V_h_* close to 0mV. This is clearly not the case with SS3. While a single data point, the SS3 structure raises concern that the expansion maybe due to an extraneous effect of salicylate and in turn raises doubt that other expanded prestin structures in salicylate are representative of the expansion due to voltage.

Previous structures of the SLC26A family members are notable for uniformly adopting an inward facing or intermediate inward facing structures. Pendrin was the only member of the family that adopted a clearly outward facing structure in the presence of both Cl^-^ and HCO ^-^ but not in the presence of either Cl^-^ or HCO ^-^ alone. Using Cl^-^ and SCN^-^, since SCN^-^ induces robust currents in prestin while preserving NLC, appears to have brought about a broadly similar result. However, there are key differences – first, the majority of structures adopted a symmetric outward (15%) structure or an asymmetric outward and inward structure (70%) in pendrin. In contrast, only a minority of structures adopted the most extreme outward facing structure (RM2) with the majority remaining in an inward facing occluded (9YAY) or intermediate state between RM2 and 9YAY. Second, the prestin structures do not adopt a cleanly outward facing form but fall short of that in pendrin particularly as it relates to the upward movement of the core domain.

We find that SCN^-^ binds to the internal anion binding site with greater affinity evidenced by the inability of salicylate to quench NLC which it does at similar concentrations of Cl^-^. There is a coincident leftward shift in *V_h_* and we speculate that the higher affinity at this site may relate to the hyperpolarizing shift in *V_h_*. Its noteworthy that the binding of SCN^-^ at the internal binding site occurs through the nitrogen and sulfur atoms that have hydrophobic and polar interactions respectively, interacting with different residues. In contrast, Cl^-^ interacts principally through the same polar interactions seen with the sulfur atom in SCN^-^. The first shells with sulfur and Cl^-^have similar maxima and minima making the hydrophobic interactions with the N atom a likely significant difference accounting for the higher SCN^-^ affinity.

What is the physiological meaning of the increasingly contracted structures? So far, no group has been able to ascertain the voltage dependent structures of prestin. Instead, the field has relied on ligand induced structures to approximate what we think would be the voltage gated structure (*11–13*). In this manner, the more contracted form of prestin that we observe with RM2, both due to a reduction the angle (ø) between monomers and an inward movement of the protein in each monomer, especially close to the inner leaflet of the membrane, could be the counterpart to the physiological voltage-induced contraction. The similarity in *z*, the unitary charge carried by each motor, in the presence of both SCN^-^ and Cl^-^ would argue that this is the case. The more contracted state we observe here and not seen previously with Cl^-^ may relate to the hyperpolarizing shift in *V_h_*. In this vein, the limited upward movement of the core domain on the gate domain in prestin with a greater contraction in the protein would also argue that it has subsumed its transporter function to its role as a motor protein

---

**Supplementary table 1.**

The table shows the RMSD values (Å) of each Tm region (Cα) of seven prestin structures after aligning the gate domains of structures with the gate domain of inward facing pendrin (7Wlk1). Also show are the RMSD values of the outward facing pendrin structure (7Wle). As is evident the core domain Tm regions show the most significant deviations with Tm1 and Tm2 showing the greatest deviations.

**Supplementary table 2.**

The table shows the statistics of cryoEM data collection and model building including refinement of models that were deposited in the PDB.

**Supplementary Figure S1**. In the presence of SCN^-^, prestin takes on a significant movement towards a contracted state that is accompanied by a rotational and modestly upward movement of the core domain on the gate domain.

Shown are atomic models of the four structures of prestin in the presence of SCN^-^ (9YAY, RM1, RM3 and RM2) and outward facing pendrin (7Wle) that have been aligned with the gate domain of inward facing pendrin (7Wlk1). **B-E.** Shows the relative motion of Tm3 and 10 that move inward and modestly upward that increases from 9YAY to RM1 to RM3 to RM2 and are compared to 7Wlk1 and 7Wle. **F-I.** Shown are the relative motion of the inner (Tm1,3,10 and 8) and outer (Tm 2, 4, 9, and 11) Tm domains of the gate domains of the four structures in the presence of SCN^-^, again showing the progressive inward and modest upward deviation of the gate domains. **J-M**. Shows the core domains that have been aligned with Wlk1 and shows minimal movements except for the cytoplasmic half of Tm 6 in RM2 that shows a significant inward deviation and opposite to that of the dolphin prestin structure in the presence of salicylate.

**Supplementary Figure S2.** There is a loss of beta sheets in the presence of SCN^-^.

**A-D.** Shows the N terminus of the protein viewed from below with the beta sheet between residues 15-20 of each monomer. In 9YAY that is lost in RM1-3. **E-H.** Shows the side view of the same segment and highlights the loss of the parallel chains in the beta sheet of 9YAY in RM1-3.

**Supplementary Figure S3.** There is no difference *Q_sp_* and *z* with Cl^-^ compared to SCN^-^. **(A,B)** Summary of *Q_sp_* and *z* across conditions, showing no detectable differences between SCN□ and Cl□ despite the shift in *V_h_*.

**Supplementary Figure S4**. In the presence of salicylate and SCN^-^, prestin reverts to an expanded state that is accompanied by a lateral and modestly downward movement of the core domain.

Shown are atomic models of the three structures of prestin in the presence of SCN^-^ and salicylate (SS1, SS2 and SS3) that have been aligned with the gate domain of 9YAY. RM2 has been included for comparison. **B-E.** Shows the relative motion of Tm3 and 10 that move outwards and modestly downward in SS1and SS2. SS3 that contains SCN^-^ at the internal binding site takes on an intermediate conformation similar to RM1. **F-I.** Shown are the relative motion of the inner (Tm1,3,10 and 8) and outer (Tm 2, 4, 9, and 11) Tm domains of the core domains of SS1 and SS2 structures in the presence of salicylate and SCN^-^, again showing the progressive outward and modest downward deviation of the core domains. **J-M**. Shows the gate domains that have been aligned with 9YAY and shows a slightly more lateral movement of the cytoplasmic half of Tm6 in SS1 and 2, compared to 9YAY.

**Supplementary Figure S5 | Cryo-EM image-processing workflow and validation of SCN□-bound prestin reconstructions.** A representative micrograph is shown from a dataset of approximately 15,000 micrographs. Movie frames were motion corrected, CTF parameters were estimated, and particles were template picked. Initial 2D classification yielded approximately 800,000 particles, with representative selected 2D class averages shown in A. B, selected particles were used for ab initio reconstruction and 3D heterogeneous classification into ten classes. Particles from selected classes were used to train Topaz for particle re-picking from the original micrographs. Each selected population was subsequently subjected to homogeneous refinement, non-uniform refinement, and local refinement. Reference-based motion correction was evaluated for all three classes and improved the resolution only for RM2; it was therefore retained only for the RM2 refinement workflow. C, this workflow yielded three final prestin reconstructions: RM1 (3.56 Å), RM3 (3.76 Å), and RM2 (3.88 Å). Gold-standard FSC curves after masking are shown below each map; the horizontal line indicates the 0.143 FSC criterion. The lower panels show the viewing-direction distributions for the final reconstructions.

**Supplementary Figure S6 | Cryo-EM image-processing workflow and final reconstructions of salicylate/SCN□-bound prestin.** Representative micrographs were obtained from approximately 11374 images. Movies were motion corrected, CTF parameters were estimated, and particles were template picked. Following 2D classification, approximately 400,000 particles were selected for ab initio reconstruction, homogeneous refinement, non-uniform refinement, and 3D classification into five classes (C1). Particles from the selected C1 classes were used for Topaz-based particle re-picking, followed by a second round of classification into five classes (C2), homogeneous refinement, non-uniform refinement, and local refinement. This workflow yielded three final cryo-EM reconstructions shown in C, with overall gold-standard FSC resolutions of 4.82 Å, 3.76 Å, and 3.91 Å, respectively, using the 0.143 criterion. Gold-standard FSC curves and angular distribution plots are shown alongside each final map.

## Methods

A cartoon summary of the cryoEm reconstructions of structures in the presence of 40mM SCN^-^and 10mM SCN^-^ with 10mM Salicylate are shown in supplementary figures S5 and S6.

### Prestin expression and purification

Full-length gerbil prestin (Meriones unguiculatus, GenBank accession AF230376), containing a C-terminal enhanced yellow fluorescent protein (EYFP) tag, was expressed and purified as described previously, with modifications. Prestin was stably expressed in HEK293 cells and membrane proteins were solubilized with digitonin. Cell pellets were resuspended in buffer containing 25 mM HEPES, pH 7.4, 200 mM NaCl, 5% glycerol, 2 mM CaCl□, DNase I and an EDTA-free protease inhibitor cocktail. Membranes were solubilized by addition of digitonin to a final concentration of 2% (w/v) and incubated for 1.5 h at 4 °C with gentle agitation. Insoluble material was removed by ultracentrifugation at 160,000 × *g* for 50 min, and the clarified supernatant was filtered through a 0.45-µm membrane. The supernatant was supplemented with 10 mM imidazole and incubated with Ni-NTA resin pre-equilibrated in buffer containing 0.02% glyco-diosgenin (GDN). The resin was sequentially washed with high-salt buffer containing 25 mM HEPES, pH 7.4, 700 mM NaCl, 5% glycerol and 0.02% GDN, followed by buffer containing 25 mM HEPES, pH 7.4, 200 mM NaCl, 30 mM imidazole, 5% glycerol and 0.02% GDN. Prestin was eluted with 25 mM HEPES, 7.4 pH, 200 mM NaCl, 250 mM imidazole, 5% glycerol and 0.02% GDN. The eluate was concentrated and further purified by size-exclusion chromatography using a Superdex 200 Increase 10/300 GL column equilibrated in 10 mM HEPES, pH 7.4, 200 mM NaCl and 0.02% GDN. Fractions corresponding to dimeric prestin were pooled and concentrated using a 100-kDa molecular-weight-cutoff centrifugal concentrator for cryo-EM sample preparation.

Two ligand conditions were prepared to investigate the conformational landscape of prestin. For the SCNO dataset, purified prestin was supplemented with 40 mM sodium thiocyanate (NaSCN) /100mM NaCl and incubated for 10 min before cryo-EM grid preparation. For the mixed salicylate/SCN□ condition, purified prestin was buffer-exchanged into a chloride-free buffer containing 10 mM HEPES, pH 7.4, 140 mM sodium gluconate and 0.02% GDN. Following buffer exchange, the sample was supplemented with 10 mM sodium salicylate and 10 mM sodium thiocyanate (NaSCN) before cryo-EM grid preparation. This buffer exchange removed residual chloride while maintaining prestin in the presence of both salicylate and SCN□. The resulting sample was used for cryo-EM grid preparation.

### Cryo-EM grid preparation and data acquisition

SCN□ and salicylate/SCN□ samples were adjusted to approximately 1 mg mlO¹ before vitrification. For both conditions, Quantifoil R1.2/1.3 gold grids (300 mesh) were glow-discharged at 23 mA for 60 s before sample application. Three sequential applications of 3 µl protein sample were made to each grid to increase particle density, with blotting between applications. Following the final application, grids were blotted for 5 s at a blot force of 0 using a Vitrobot Mark IV (Thermo Fisher Scientific) maintained at 8 °C and 100% relative humidity and immediately plunge-frozen in liquid ethane. Vitrified grids were stored in liquid nitrogen until cryo-EM data collection.

Cryo-EM datasets were collected using a Titan Krios transmission electron microscope operating at 300 kV, equipped with a Gatan K3 direct electron detector and a 20-eV energy-filter slit. Movies were acquired in super-resolution mode using SerialEM. The established SCN and Salicylate/SCN mixed datasets were used a nominal magnification of 81,000×, corresponding to a super-resolution pixel size of 0.534 Å, with a cumulative electron exposure of approximately 54 e□ Å□² and a nominal defocus range of −2.0 to −2.5 µm. For the 40 mM SCN□ dataset, 15,779 movies were collected. For the 10 mM salicylate/10 mM SCN□ dataset, 11,374 movies were collected.

### Cryo-EM image processing and conformational classification

Cryo-EM data were processed using CryoSPARC v4.7.1. Movie frames were subjected to patch-based motion correction, and contrast transfer function (CTF) parameters were estimated using Patch CTF estimation. Micrographs with poor CTF fits or excessive motion were excluded from subsequent processing.

For the SCN□ dataset, initial particle picking yielded approximately 2 million particles, which were extracted and subjected to iterative two-dimensional (2D) classification. Classes corresponding to false-positive picks, contaminants, poorly resolved particles, or classes lacking recognizable protein features were excluded. Approximately 300,000 selected particles displaying well-defined prestin features were subjected to homogeneous refinement. These particles were subsequently subjected to three-dimensional (3D) classification into ten classes. The least-populated class retaining clearly recognizable prestin features was selected as a high-confidence particle set for training the Topaz neural-network particle picker. The trained Topaz model was then applied to the dataset for particle repicking. Topaz-picked particles were extracted and subjected to further 2D classification and particle cleaning, followed by ab initio reconstruction, homogeneous refinement, non-uniform refinement, 3D classification and local refinement. The resulting particle populations were independently refined to obtain the three SCN□-bound conformations: RM1 from 157,912 particles, RM2 from 75,736 particles and RM3 from 71,752 particles, at global resolutions of 3.56 Å, 3.88 Å and 3.76 Å, respectively.

For the mixed salicylate/SCN□ dataset, particle picking and extraction initially yielded 4,628,413 particles, which were subjected to 2D classification into 100 classes. Classes displaying well-defined protein features were selected, retaining 1,802,296 particles. A second round of 2D classification into 50 classes, followed by removal of poorly resolved classes, retained 1,567,051 particles. Following additional particle selection, 1,377,300 particles were subjected to ab initio reconstruction into four classes, followed by homogeneous refinement. To further resolve conformational heterogeneity, selected particles were subjected to 3D classification into ten classes. A particle population representing a distinct prestin conformation was selected as a high-confidence training set for the Topaz neural-network particle picker. The trained Topaz model was subsequently applied to the dataset for particle repicking. Topaz-picked particles were extracted and subjected to further particle cleaning, followed by ab initio reconstruction, homogeneous refinement, non-uniform refinement and additional 3D classification. The resulting conformational populations were independently refined to obtain SS1 from 88,936 particles, SS2 from 94,630 particles and SS3 from 183,608 particles, at global resolutions of 4.88 Å, 3.76 Å and 3.90 Å, respectively.

Final map resolutions were determined using the gold-standard Fourier shell correlation criterion at FSC = 0.143, and local-resolution estimation was performed. Final particle numbers, map resolutions and map-sharpening B factors for the individual reconstructions are provided in the cryo-EM data collection, refinement and validation statistics table.

### Atomic model building and refinement

The previously determined gerbil prestin structure (PDB 7SUN) was used as the starting model for atomic model building of the RM1–RM3 and SS1–SS3 cryo-EM reconstructions. The 7SUN model was rigid-body fitted into each cryo-EM density map using UCSF Chimera/ChimeraX. The fitted models were initially subjected to real-space refinement in Phenix and subsequently inspected and manually adjusted in Coot by examining the complete protomer chains. Particular attention was given to regions exhibiting conformational differences among the six reconstructions, including the transmembrane helices, the relative positions of the core and gate domains, and the cytoplasmic STAS domains. Each model was refined through iterative rounds of manual model building in Coot and real-space refinement in Phenix using secondary-structure and geometry restraints. Model geometry and stereochemical quality were evaluated using MolProbity and the validation tools implemented in Phenix. Across the six final models, bond-length root-mean-square deviations (RMSDs) ranged from 0.003 to 0.004 Å and bond-angle RMSDs from 0.597° to 0.702°. MolProbity scores ranged from 1.69 to 2.29 and clash scores from 4.51 to 9.58. Ramachandran analysis showed 90.06–93.35% of residues in favored regions, with 0–0.30% Ramachandran outliers. Rotamer outliers ranged from 0 to 3.12%. Detailed refinement and validation statistics for each reconstruction are provided in the cryo-EM data collection, refinement and validation statistics supplementary table. For the SCN□-bound RM structures (RM1–RM3), density at the internal anion-binding site was inspected for bound SCN□, and SCN□ was modeled where supported by the experimental density and local binding-site environment. For the mixed salicylate/SCN□ structures (SS1–SS3), ligand density at the internal anion-binding site was evaluated independently in each reconstruction. Salicylate/SCN-was modeled at the internal site supported by the experimental density and local chemical environment and was refined together with the protein model in Phenix.

### Potential hydrogen bond modeling

Potential hydrogen bonds in the N-terminal β-sheet region of the prestin structures were modelled with UCSF ChimeraX. Residues 16–20 in both chains A and B were selected and analyzed their hydrogen-bond interactions determined using the hbonds command with the restrict both options. The default ChimeraX hydrogen-bond detection criteria, which consider donor–acceptor distances and hydrogen-bond geometry were used. No additional distance cutoff was specified.

### Electrophysiological recording of prestin NLC

Prestin stably expressed in a tetracycline inducible HEK line was recorded in a whole-cell configuration at room temperature using an Axon 200B amplifier (Molecular Devices, Sunnyvale, CA), as described previously (*18, 23*). Whole-cell recordings were performed 24–72 h after tetracycline induction.

To minimize extracellular Cl⁻, the bath solution was gluconate-based, retaining only a small amount of Cl⁻ (0.2 mM CaCl) to support recording stability. The bath solution contained (in mM): 100 NaGluconate, 2 MgGluconate, 0.2 CaCl_2_, 10 Hepes, pH 7.2, 300 mOsm. A series of pipette solutions with different concentrations of Cl^-^ or SCN^-^ was used. The base pipette solution contained (in mM): 20 CsGluconate, 5 EGTA, 2 MgGluconate, 0.2 CaCl_2_, 10 Hepes, with NaGluconate adjusted to maintain a total of 100 mM combined NaGluconate + test anion. For the Cl^-^ series, NaCl was added to base solution at 0.5, 1, 5, 10, 40, 100 mM (with a corresponding reduction of NaGluconate). For the SCN^-^ series, NaSCN was added to base solution at 0.5, 1, 5, 10, 40, 100 mM (with a corresponding reduction of NaGluconate). Osmolarity was adjusted to 300 ± 2 mOsm with dextrose and all solutions were buffered to pH 7.2.

NLC was measured using a continuous, high-resolution two-sine voltage command. Command delivery and data acquisition were controlled with jClamp (Scisoft, Ridgefield, CT) via a Digidata 1322A interface (Axon Instruments). Capacitance–voltage data were fit with the first derivative of a two-state Boltzmann function:

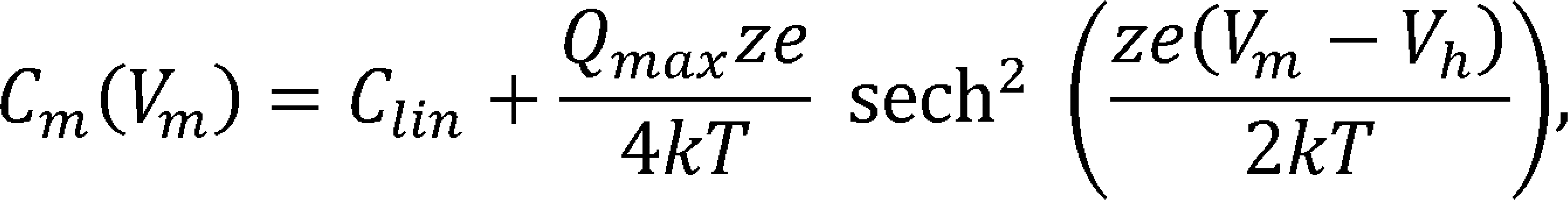

where *Q_mox_* is the maximum nonlinear charge transfer, *V_h_* is the voltage at peak capacitance (or half-maximal nonlinear charge transfer), *V_m_* is membrane potential, *C_Iİn_* is the linear capacitance, *z* is the apparent unitary charge (voltage sensitivity), *e* is the elementary charge, *k* is the Boltzmann constant, and *T* is absolute temperature. Specific charge density was computed as:

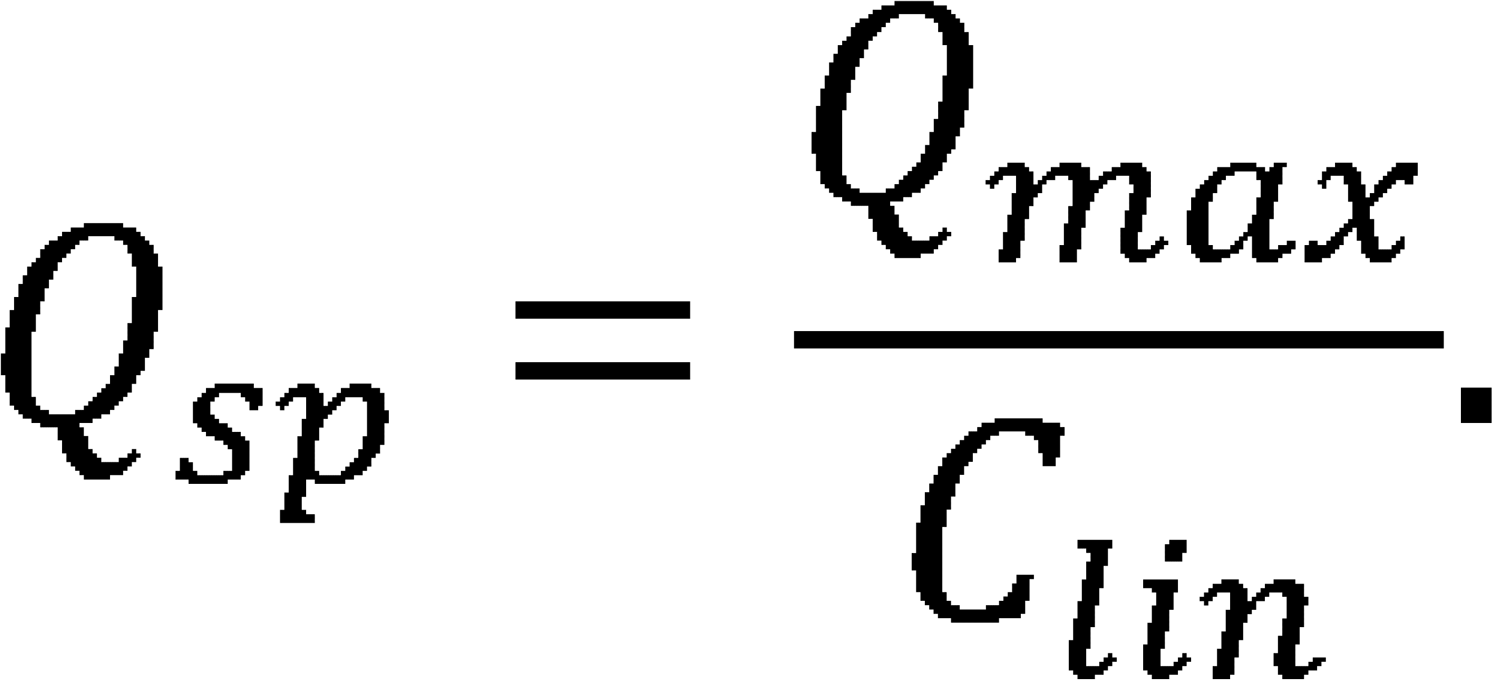

### Restrained molecular dynamics simulation setup

The model systems were prepared using the CHARMM-GUI workflow (*24, 25*) and continued with 100 ns of restrained MD. These trajectories are distinct from unrestrained production runs. Simulations were performed with GROMACS 2024.6 (*26, 27*) at 296 K and 1 bar.Harmonic positional restraints were applied to protein backbone atomswith a force constant of 50 kJ mol-1 nm-2; positional restraints on protein side chains and lipids, and dihedral restraints, were set to zero.

Temperature was maintained at 296 K using the stochastic velocity-rescaling thermostat (*28*), with separate coupling groups for solute (SOLU), membrane (MEMB), and solvent (SOLV) and a 1 ps coupling time for each group.

Semi-isotropic pressure coupling maintained 1 bar with a 5 ps coupling time and a compressibility of 4.5 x 10^-5^ bar^-1^. The RM1-RM3 simulations used the Parrinello-Rahman barostat (*29*), whereas the SS and 9YAY simulations used the stochastic cell-rescaling (C-rescale) barostat (*30*).

Long-range electrostatics were treated with PME using a 1.2 nm Coulomb cutoff (*31*). Lennard-Jones interactions used a force-switch modifier, switched from 1.0 nm to zero at 1.2 nm. A Verlet cutoff scheme with a 1.2 nm neighbor-list cutoff was used, and bonds to hydrogen atoms were constrained with LINCS (*32*).

#### Area Calculation

Protein cross-sectional area profiles were calculated for each trajectoryframe using a periodic two-dimensional Voronoi tessellation (*21*). Trajectories were centered on the bilayer center of mass, and the system was divided into slices perpendicular to the membrane normal (the z-axis) at 0.5 Å intervals. Within each slice, atomic positions were assigned to Voronoi cells; the areas of cells belonging to protein atoms were summed to obtain the area profile A(t,z). The maximum cross-sectional area for each frame was taken as max_z A(t,z) and used for the statistical comparisons.

#### Statistical analysis

For statistical comparisons, maximum cross-sectional area trajectories from eight systems (SS1-SS3, RM1-RM3, 7SUN, and 9YAY) were analyzed over the final 50% of each trajectory, with frames sampled every 0.01 ns. Temporal autocorrelation was estimated from the integrated autocorrelation time using Geyer’s initial-positive sequence (*33*).

Comparisons used three complementary approaches: Welch’s t-tests with correlation-adjusted effective sample sizes (*34*), Welch’s t-tests on data subsampled at twice the maximum estimated correlation time (3.75 ns), and a moving-block bootstrap with 3.75 ns blocks and 20,000 resamples (*35, 36*). Each system was compared with 9YAY, and all 28 system pairs were also tested; Holm correction was applied separately within each comparison family and method (*37*). RM2 had the lowest mean maximum cross-sectional area (3,131.0 A^2) and differed significantly from each of the other seven systems in all three autocorrelation-aware pairwise analyses (Holm-adjusted p < 0.05). In direct comparisons, RM2’s mean was 69.3 A^2 lower than RM1 and 65.0 A^2 lower than RM3; these differences remained significant after Holm correction across the 28 pairwise tests for each method (adjusted p <= 0.0082).

