## Supplementary figures and images for "SCN^-^ induces prestin contraction through reduced intermonomer angles and a modified transporter like movement of each monomer"

### Supplementary Figure 1

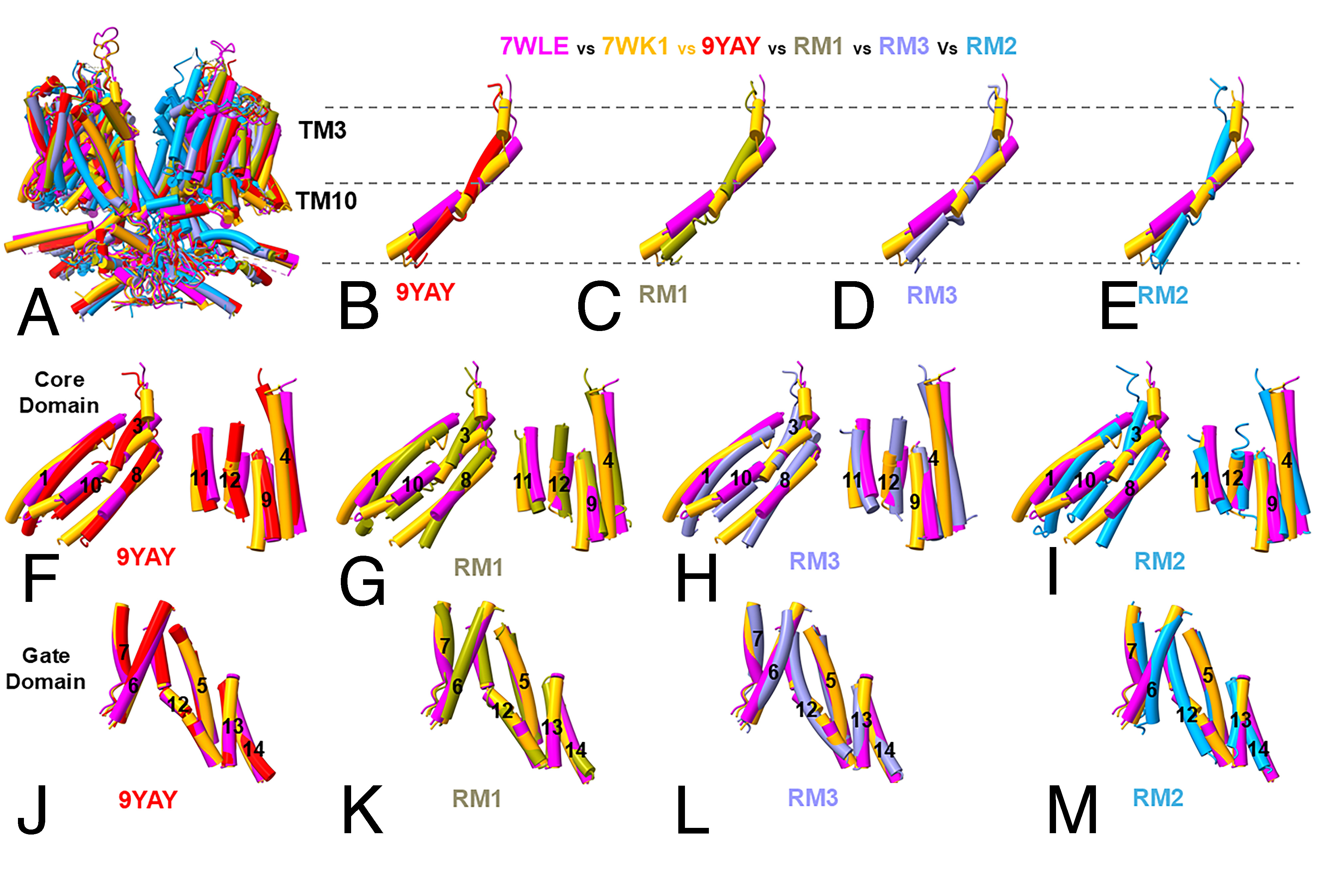

### Supplementary Figure 2

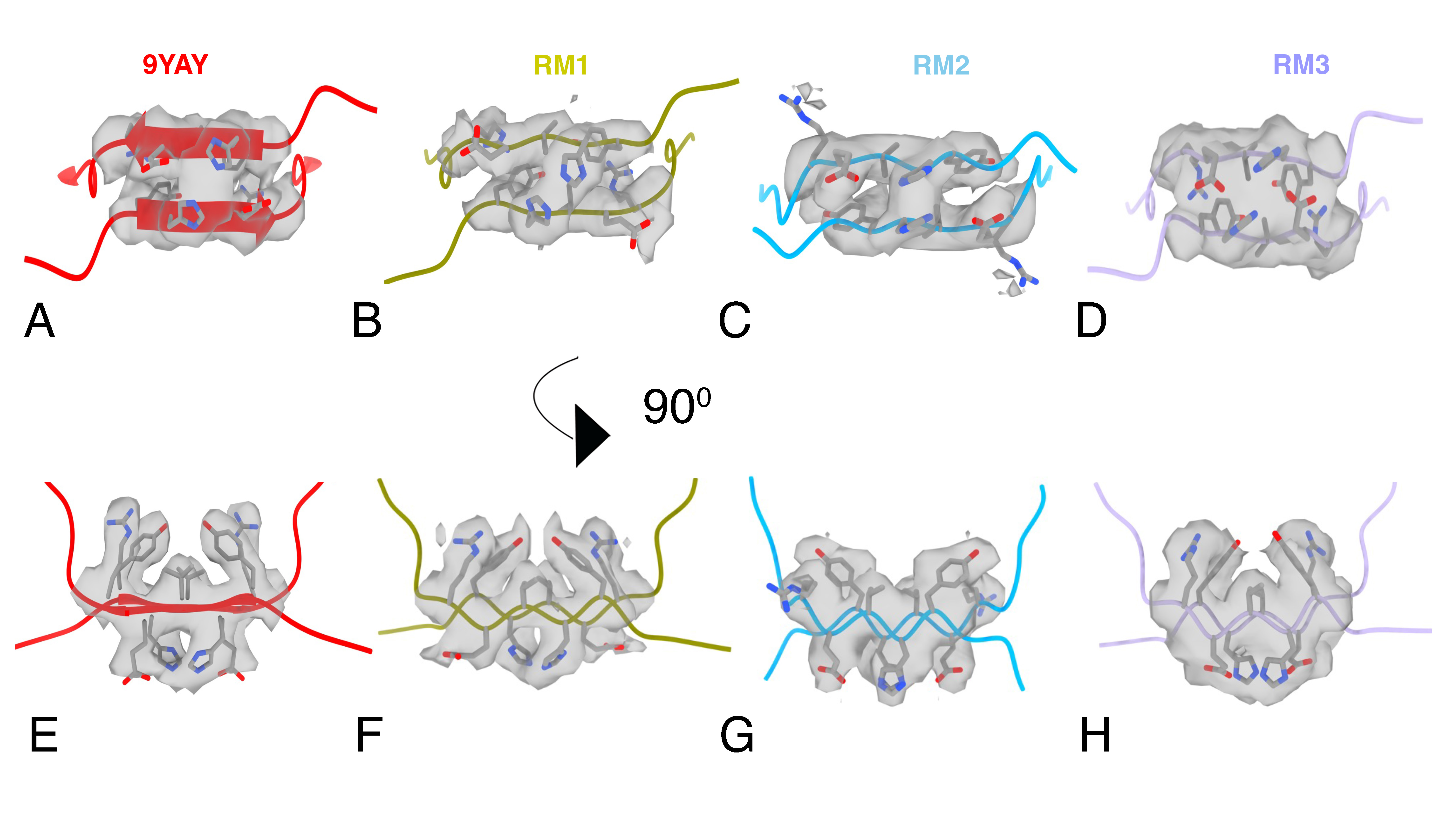

### Supplementary Figure 3

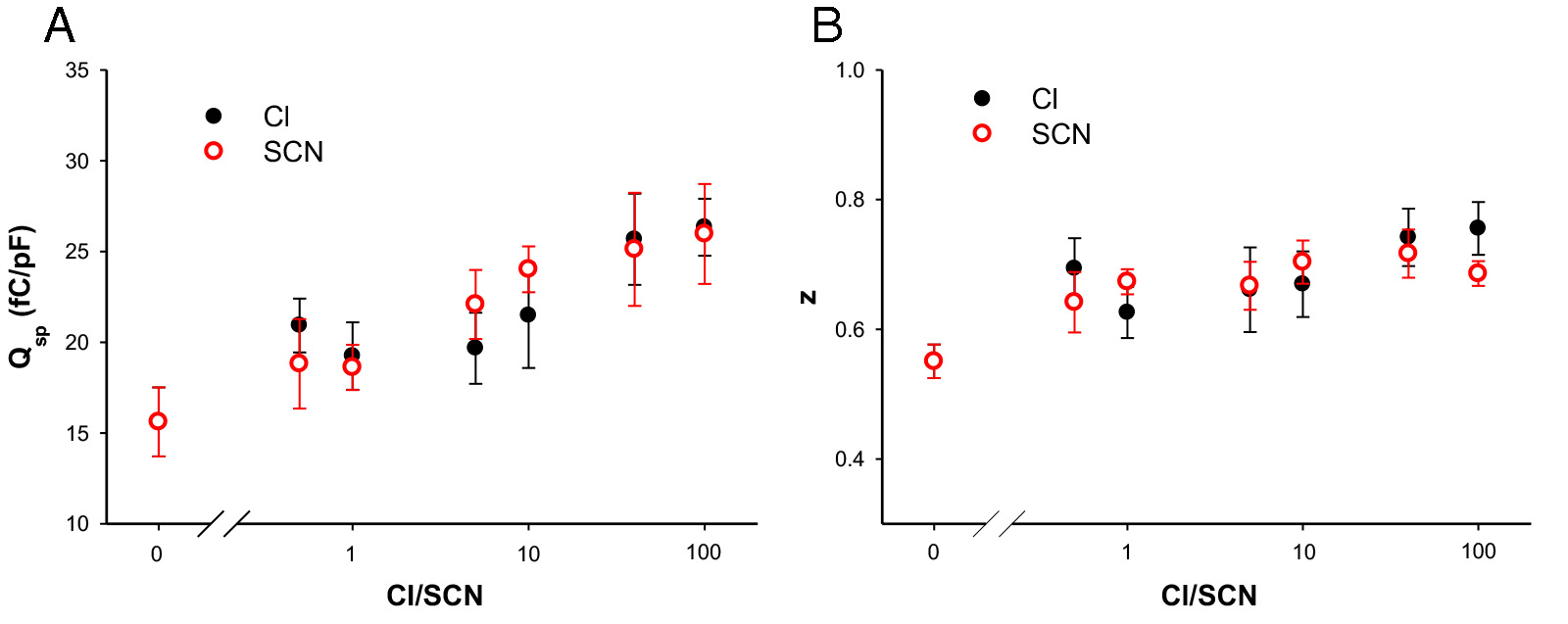

### Supplementary Figure 4

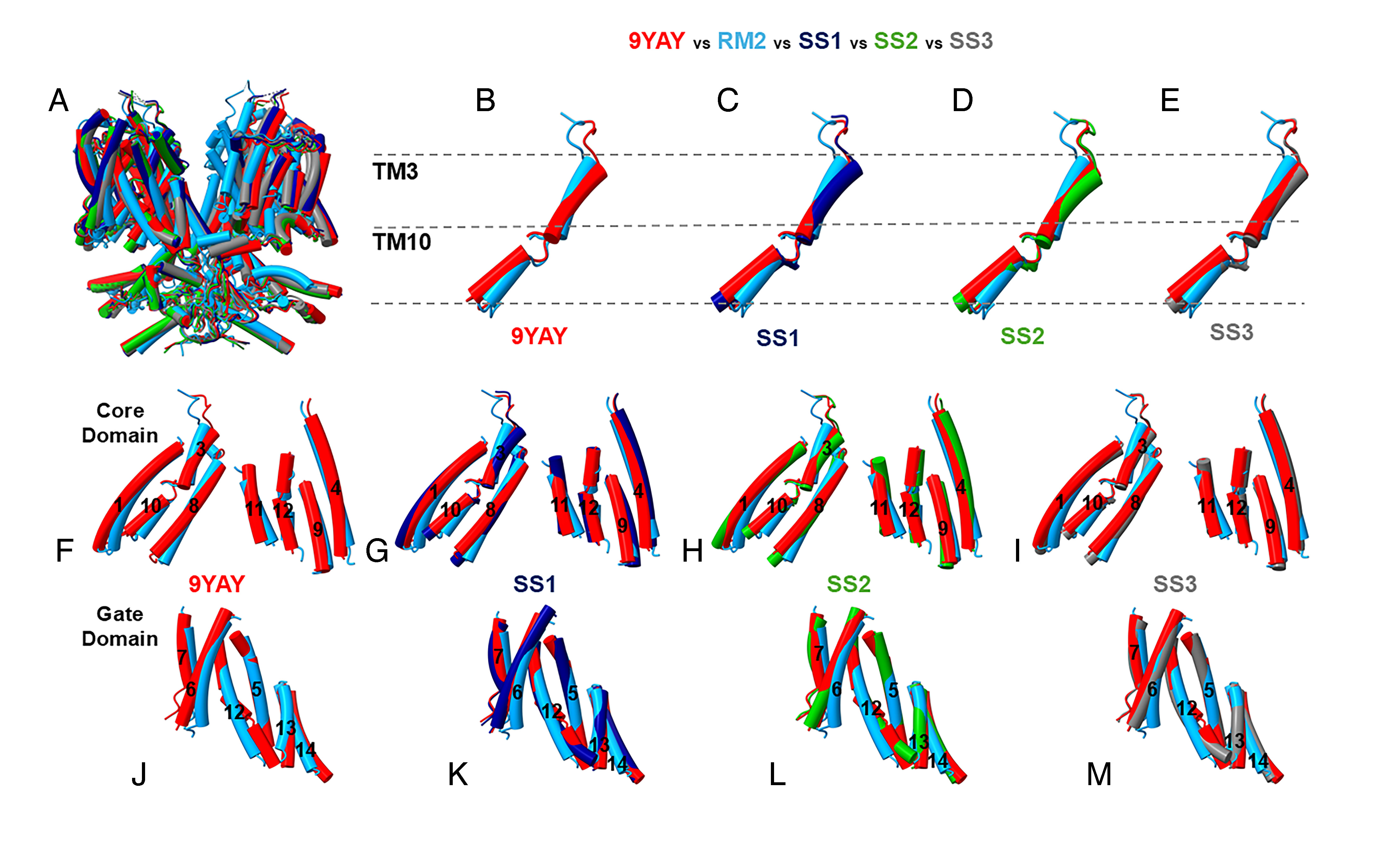

### Supplementary Figure 5

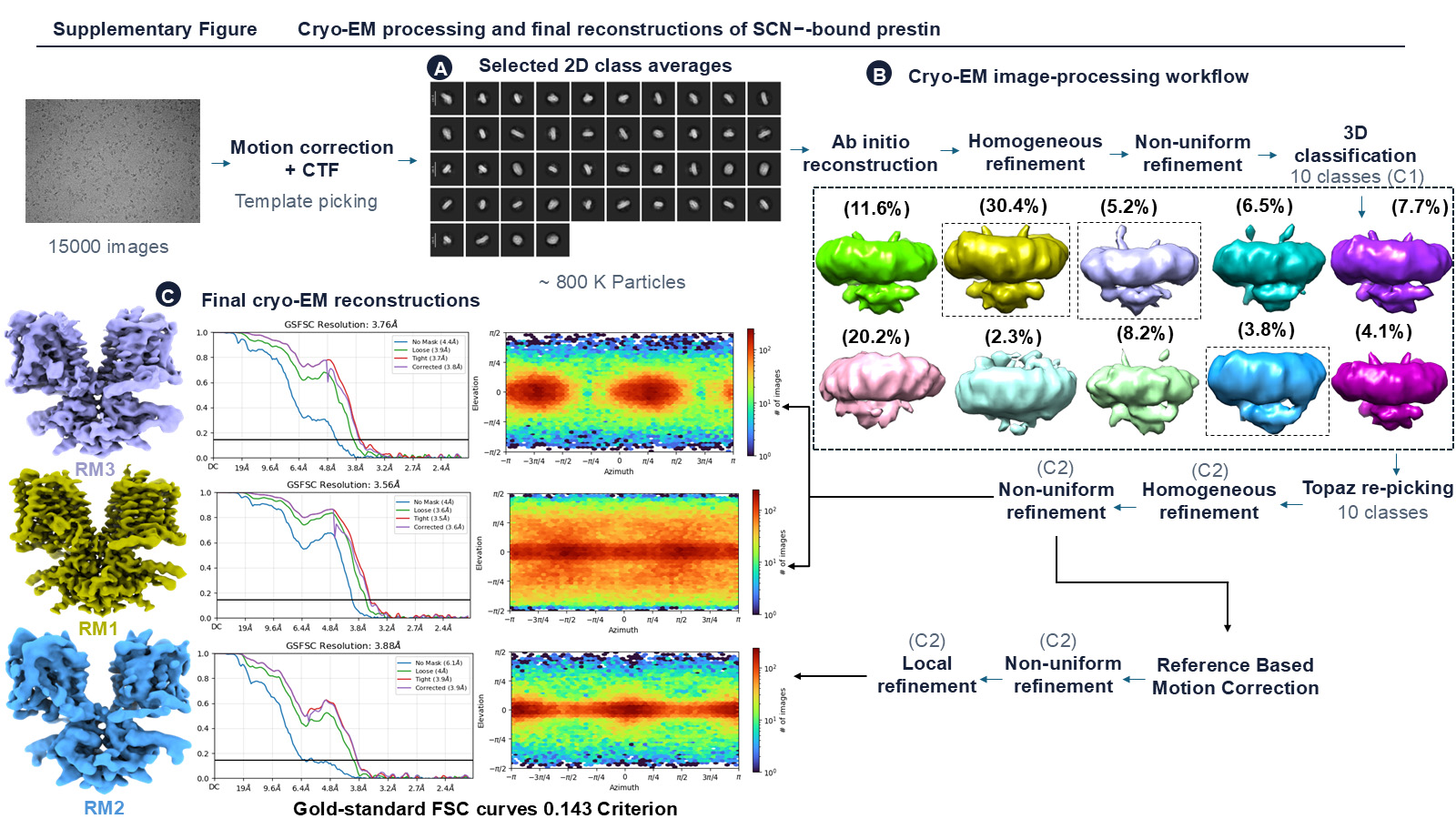

### Supplementary Figure 6

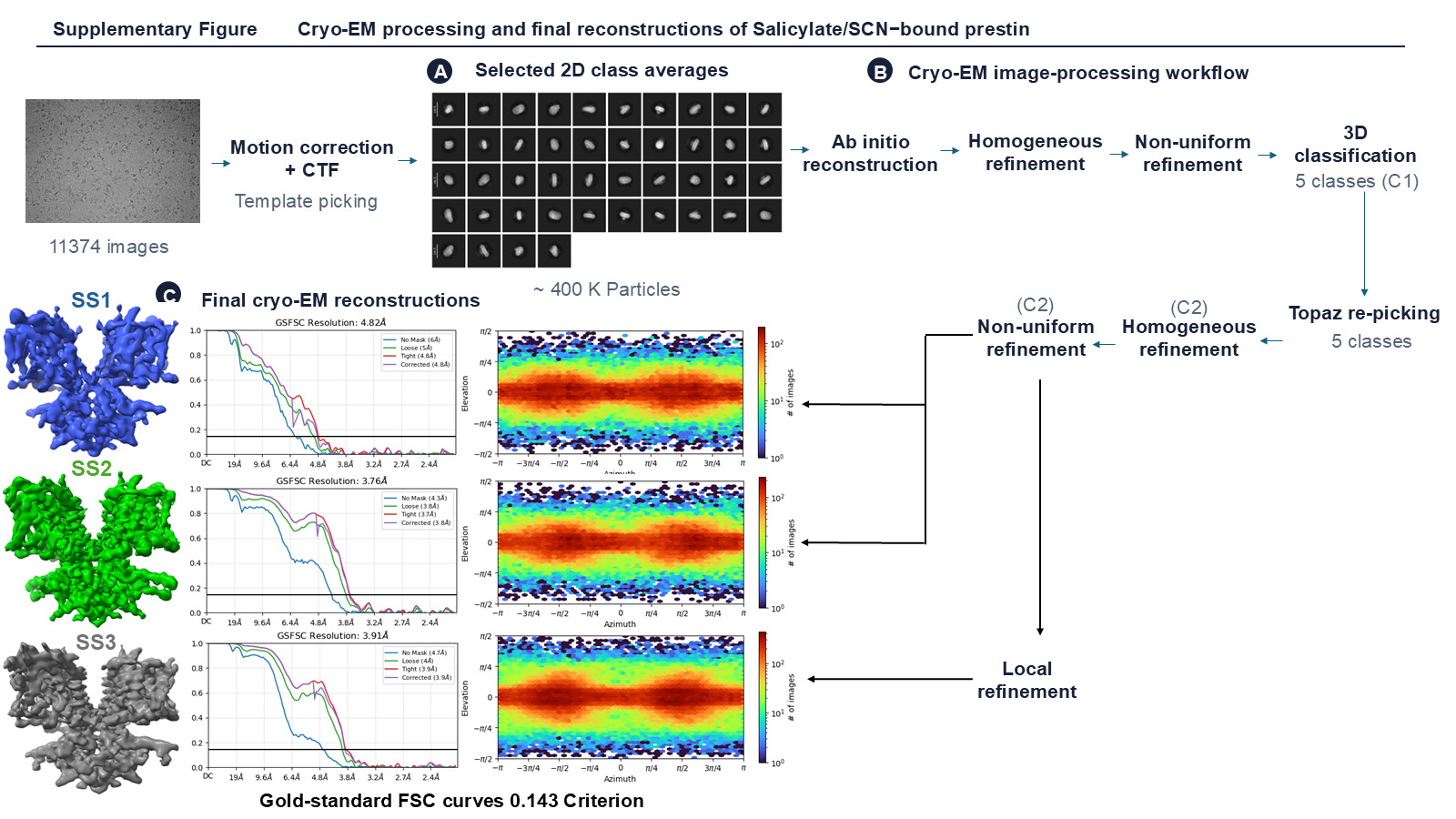
