## Supplementary Table 1 for "SCN^-^ induces prestin contraction through reduced intermonomer angles and a modified transporter like movement of each monomer"

| **Helix** | **Domain** | **Prestin residues** | **Pendrin residues** | **Cα pairs** | **9YAY** | **RM1** | **RM2** | **RM3** | **SS1** | **SS2** | **SS3** | **7Wle** |
| --- | --- | --- | --- | --- | --- | --- | --- | --- | --- | --- | --- | --- |
| **TM1** | **Core** | **75–106** | **85–116** | **32** | **9.393** | **9.342** | **10.266** | **9.704** | **9.268** | **9.280** | **10.065** | **7.707** |
| **TM2** | **Core** | **109–127** | **119–137** | **19** | **11.214** | **11.228** | **10.798** | **11.181** | **11.505** | **11.322** | **11.338** | **3.310** |
| **TM3** | **Core** | **137–151** | **141–155** | **15** | **5.449** | **5.571** | **6.417** | **5.491** | **5.109** | **4.155** | **4.756** | **8.816** |
| **TM4** | **Core** | **167–196** | **177–206** | **30** | **3.936** | **4.308** | **4.991** | **4.200** | **4.600** | **3.504** | **3.972** | **5.958** |
| **TM5** | **Gate** | **206–230** | **216–240** | **25** | **1.191** | **1.258** | **1.507** | **1.238** | **1.032** | **0.856** | **0.917** | **0.763** |
| **TM6** | **Gate** | **258–283** | **268–293** | **26** | **0.833** | **0.912** | **2.793** | **1.419** | **1.983** | **1.069** | **1.025** | **0.713** |
| **TM7** | **Gate** | **288–306** | **298–316** | **19** | **0.789** | **0.900** | **3.822** | **1.592** | **0.842** | **1.200** | **1.596** | **0.465** |
| **TM8** | **Core** | **344–366** | **352–374** | **23** | **5.908** | **6.199** | **6.419** | **6.076** | **5.819** | **5.276** | **5.621** | **8.401** |
| **TM9** | **Core** | **370–388** | **380–398** | **19** | **3.811** | **3.934** | **4.128** | **3.609** | **4.443** | **3.048** | **3.800** | **3.762** |
| **TM10** | **Core** | **397–407** | **406–416** | **11** | **5.683** | **5.906** | **7.490** | **6.084** | **5.465** | **5.155** | **5.981** | **9.024** |
| **TM11** | **Core** | **412–427** | **422–437** | **16** | **3.117** | **2.951** | **4.224** | **3.629** | **2.842** | **2.604** | **2.734** | **4.154** |
| **TM12** | **Gate** | **436–463** | **446–473** | **28** | **1.037** | **1.117** | **3.085** | **1.440** | **1.051** | **0.955** | **1.562** | **0.781** |
| **TM13** | **Gate** | **465–481** | **475–491** | **17** | **0.948** | **0.949** | **1.439** | **1.197** | **0.726** | **1.009** | **0.985** | **0.523** |
| **TM14** | **Gate** | **483–505** | **493–515** | **23** | **1.652** | **1.629** | **3.139** | **1.710** | **1.695** | **1.338** | **1.717** | **0.743** |
