## Supplementary Table 2 for "SCN^-^ induces prestin contraction through reduced intermonomer angles and a modified transporter like movement of each monomer"

Supplementary Table 2: Statistics and refinement

| **Parameter** | **RM1** | **RM2** | **RM3** | **SS1** | **SS2** | **SS3** |
| --- | --- | --- | --- | --- | --- | --- |
| **Image processing** | | | | | | |
| Microscope | Titan Krios | Titan Krios | Titan Krios | Titan Krios | Titan Krios | Titan Krios |
| Voltage (kV) | 300 | 300 | 300 | 300 | 300 | 300 |
| Camera | Gatan K3 | Gatan K3 | Gatan K3 | Gatan K3 | Gatan K3 | Gatan K3 |
| Magnification | 81,000 | 81,000 | 81,000 | 81,000 | 81,000 | 81,000 |
| Electron exposure (e⁻ Å⁻²) | 54 | 54 | 54 | 54 | 54 | 54 |
| Defocus range (µm) | −2.0 to −2.5 | −2.0 to −2.5 | −2.0 to −2.5 | −2.0 to −2.5 | −2.0 to −2.5 | −2.0 to −2.5 |
| Pixel size (Å) | 0.534 | 0.534 | 0.534 | 0.534 | 0.534 | 0.534 |
| Symmetry imposed | C2 | C2 | C2 | C2 | C2 | C2 |
| Micrographs | 15,779 | | | 11,374 | | |
| Final particle images | 157,912 | 75,736 | 71,752 | 88,936 | 94,630 | 183,608 |
| Map resolution (Å) | 3.56 | 3.88 | 3.76 | 4.82 | 3.76 | 3.90 |
| FSC threshold | 0.143 | 0.143 | 0.143 | 0.143 | 0.143 | 0.143 |
| Map-sharpening B factor (Å²) | **133.4** | **89.2** | **121.3** | **276.1** | **125.9** | **128.9** |
| **Model refinement** | | | | | | |
| Bond RMSD (Å) | 0.004 | 0.003 | 0.003 | 0.003 | 0.004 | 0.003 |
| Angle RMSD (°) | 0.661 | 0.702 | 0.597 | 0.658 | 0.636 | 0.624 |
| MolProbity score | 2.11 | 2.29 | 1.69 | 1.90 | 2.17 | 2.28 |
| Clashscore | 6.26 | 9.58 | 4.51 | 7.53 | 8.77 | 8.68 |
| Rotamer outliers (%) | 2.35 | 2.08 | 0.69 | 0.00 | 1.91 | 3.12 |
| Ramachandran favored (%) | 92.00 | 90.06 | 92.53 | 91.93 | 92.00 | 93.35 |
| Ramachandran allowed (%) | 7.92 | 9.64 | 7.40 | 8.07 | 8.00 | 6.65 |
| Ramachandran outliers (%) | 0.07 | 0.30 | 0.07 | 0.00 | 0.00 | 0.00 |
